# A humanized Aβ mouse model reveals *E4*-dependent cognitive impairments, microglial activation, and cerebrovascular dysfunction

**DOI:** 10.64898/2026.07.28.740806

**Authors:** Skylyn J Ferguson, Claudia Pelayo, Siena Stueland, Kennedy Krajack, Kiara Gardley, Robert Herbert, Caitlin Tafoya, Alex Famiano, Abigail E Cullen, Naly Setthavongsack, Randall L. Woltjer, Ashley E Walker

## Abstract

Apolipoprotein *E4* (*E4*) increases the risk of Alzheimer’s disease (AD) by up to 12-fold. However, understanding of the mechanisms underlying this increased risk has been limited by a lack of preclinical models that accurately reflect the effects of *E4* in the presence of humanized non-mutant amyloid-β precursor protein (*hAβPP*). Therefore, we studied novel humanized *APOE* and *hAβPP* mice to investigate the contributions of the *E4* genotype to cognitive, inflammatory, and vascular dysfunction, specifically comparing male and female *E3/hAβPP* and *E4/hAβPP* mice. *E4/hAβPP* mice exhibited impaired nest-building behavior and novel object recognition compared with *E3/hAβPP* mice. Microglial content was higher in *E4/hAβPP* mice, whereas astrocyte content was not different across groups. *E4/hAβPP* mice had greater carotid and cerebral artery stiffness, and higher collagen I content in cerebral arteries than *E3/hAβPP* mice. Under static pressure, cerebral artery endothelium-dependent and endothelium-independent vasodilation were similar across genotypes. However, high pulse pressure selectively impaired cerebral artery endothelial function in *E4/hAβPP* mice, with the greatest impairment observed in females. The *E4/hAβPP* mice also exhibited higher cortical expression of *Nox2* and *Sod1* and elevated cerebral artery *Il1b* expression. As such, *E4/hAβPP* mice exhibit convergent cognitive, inflammatory, and vascular abnormalities that recapitulate several features of AD. Elevated pulse pressure revealed an *E4*-dependent vulnerability of the cerebral vasculature, suggesting that vascular stress may be an important contributor to disease risk. Together, our findings support the use of the *APOExhAβPP* model to investigate the mechanisms by which *E4* promotes vascular dysfunction, neuroinflammation, and cognitive impairment in AD.

## Introduction

Despite the extensive research conducted to cure or prevent Alzheimer’s disease (AD), the mechanisms that initiate and drive AD pathogenesis remain incompletely understood. The therapeutic development for AD is hindered by the limited translational relevance of most preclinical models used to study AD. The most commonly used models rely on familial AD mutations such as those in the amyloid-β protein precursor (*AβPP*) and the presenilin 1 (*PS1*) genes^1,2^. The familial AD mutations only account for less than 1% of human AD cases^3^ and neglect the genetic differences that contribute to late-onset AD (LOAD).

Out of all the genetic risk factors for LOAD, the apolipoprotein ε4 (*E4)* genotype is the strongest^4^. *E4* accelerates amyloid deposition, alters lipid metabolism, disrupts synaptic function, and contributes to neuroinflammation^5^. In contrast, the apolipoprotein ε3 (*E3)* is considered neutral to an individual’s AD risk, while the apolipoprotein ε2 is considered protective^6,7^. Since *E4* influences multiple upstream pathways, such as Aβ clearance^5^, microglial activation^8^, and neuroinflammation responses^9^, models that incorporate human *APOE* isoforms are more physiologically relevant to study for the mechanistic translation between pre-clinical models and human AD patients. However, research studies involving *APOE*-targeted mouse models have been limited by the absence of humanized Aβ, or they rely on aggressive familial AD mutations, such as the 5xFAD and *AβPP*x*PS1* mouse models, that overshadow *APOE*-specific effects ^1,2,10,11^. To address these limitations, we sought to understand the impact of *APOE* genotype in the context of humanized, non-mutant *AβPP*. To do so, we studied the *E4xhAβPP* and *E3xhAβPP* strains recently developed by MODEL-AD. These models combine humanized *APOE* alleles with a knock-in humanized *AβPP* sequence designed to mimic the endogenous Aβ production by non-mutant *AβPP*. To our knowledge, this is the first report of vascular outcomes from these models, which provide a deeper understanding of *APOE*-dependent mechanisms of AD.

The *E4* genotype is not only the strongest genetic risk factor for LOAD, but is also closely linked to cerebrovascular dysfunction^12,13^. *E4* carriers show a greater age-related decline in cerebral blood flow (CBF) than the more common *E3* genotype ^14^, and insufficient CBF to activated brain regions contributes to neurodegeneration and cognitive impairment ^15^. Under healthy conditions, endothelial nitric oxide (NO) is produced to promote vasodilation, a process known as endothelium-dependent dilation^15^. However, *E4* is associated with impaired endothelial signaling, heightened oxidative stress, and increased production of reactive oxygen species, all of which diminish NO bioavailability and disrupt vascular regulation^16–19^. Superoxide, in particular, reacts rapidly with NO to form peroxynitrite, further reducing NO signaling and damaging endothelial function^20^. Deficits in NO mediated dilation and increased oxidative stress are thought to contribute to the accelerated cognitive decline observed in *E4* carriers^21–23^, yet the precise mechanisms linking *E4*, vascular dysfunction, and heightened LOAD risk remain incompletely understood.

Advancing age is accompanied by increases in pulsatile pressure, which impose greater mechanical stretch on cerebral arteries and arterioles and impair endothelial function^24,25^. Higher pulse pressure reduces NO signaling, disrupts CBF regulation, and is associated with poorer executive function in humans ^15,26^. Excessive mechanical stretch promotes vascular remodeling, elevates inflammatory signaling, and increases reactive oxygen species production, including superoxide, which further compromises endothelial health^27,28^. A highly oxidative and inflammatory environment not only diminishes NO bioavailability but also contributes to cerebral arterial stiffness by negatively impacting the extracellular matrix environment, exacerbating cognitive decline^24,29–31^. *E4* carriers have stronger associations between elevated arterial stiffness and systolic blood pressure with cognitive impairment^12,32–34^ but the interaction between *E4* and high pulse pressure has not been explored.

This study was designed to determine how the *E4* genotype impacts cognitive function, neuroinflammation, oxidative stress, and cerebral artery function in a non-mutant *hAβPP* model. We also built these studies to determine the interaction effect of the *E4* genotype and high pulse pressure on cerebral vascular function. We hypothesized that the *E4* mice would have greater memory impairments, neuroinflammation, arterial stiffness, oxidative stress, and cerebral artery endothelial dysfunction compared to the *E3* mice. We further hypothesized that the cerebral arteries from *E4* mice would have greater impairments to endothelial function after exposure to high pulse pressure than cerebral arteries from *E3* mice. To test our hypotheses, we studied young homozygous *E3xhAβPP* and *E4xhAβPP* mice. We tested for potential sex differences because of the higher prevalence of AD in females^35^. In these mice, we assessed learning and memory through novel object recognition and instinctual behavior with the nest building task. The large artery stiffness of the mice was tested using *in vivo* aortic pulse wave velocity. We then collected their cerebral arteries to assess the endothelium-dependent dilation after exposure to static pressure, and low and high pulse pressures. We also collected arteries to measure both cerebral artery and carotid artery stiffness. We assessed potential mechanisms for our functional outcomes through immunofluorescence, immunohistochemistry, gene expression, and protein expression.

## Methods

### Animals

Male and Female mice homozygous for either *E4* or *E3* and humanized amyloid-precursor protein (*E4xhAβPP* and *E3xhAβPP*) were obtained from the Jackson Laboratory and studied at 6 months of age (strain #’s: 034455 and 034415). All mice were fed a phytoestrogen-free diet (Envigo, Teklad Global Soy Protein-Free Extruded Rodent Diet, 2920X) with food and water provided ad libitum and housed in a facility with a 12/12-h light-dark cycle at 24°C. Mice were euthanized by exsanguination under isoflurane (2% in 100% O_2_) anesthesia and perfused with heparin saline immediately before ex vivo vascular reactivity experiments. All animal procedures conform to the *Guide to the Care and Use of Laboratory Animals* (8^th^ edition, revised 2011) and were approved by the Institutional Animal Care and Use Committee at the University of Oregon.

### Aortic Stiffness

Aortic stiffness was measured *in vivo* by pulse wave velocity (PWV) using Indus Doppler Flow Velocity System (Webster, Texas) with a previously established protocol from our lab ^24,36^. Two researchers analyzed each Doppler recording and inter-observer confidence intervals were measured, and PWV measurements with confidence intervals over 10% were excluded.

### Cognitive Testing

The nest building test was used to assess instinctual cognitive behavior. Mice were singly housed overnight in a cage with food, water and a condensed cotton nestlet. After the mice completed their dark cycle, a researcher, blinded to the genotypes and sex of the mice, scored the nests on a scale of 0-5 based on the quality of their nest ^37^. Accelerating rotarod was used to assess motor function in the mice as a potential confounding variable for other cognitive tests^38^. On the acclimatization day, the mouse had to stay on the rotating rod for 90 seconds at a constant speed of 4 rpm. On the second day of the rotarod test, the mice ran three separate trials on the rod as it accelerated from 4 rpm to 40 rpm. The time at which the mouse fell off the rod or rotated around the rod twice was recorded and did not exceed 300 seconds.

An open field test was used to assess mouse anxiety levels. The open field test consisted of mice exploring a 40 cm^2^ square arena with a white matte floor and walls for 10 minutes. The exploration of the mice was recorded using Ethovision XT12 software. In the middle of the arena, a 20 cm^2^ square was marked with Ethovision Software to denote the inner area of the arena. The exploration criterion was defined as the amount of time the center of the mouse’s body was inside the inner square area. Mice who spent more time exploring the inner square were considered to have lower levels of anxiety.

One day after the open field test was performed, mice underwent the Novel Object behavioral test for spatial and short-term memory testing. Mice were placed in the same arena used for open field with two of the same objects. Mice explored the two objects for 10 minutes. 1 hour later, the mice were placed in the arena again with one familiar object and one novel object and explored for 10 minutes. The side of the novel object was alternated between mice to avoid any biases. Ethovision Software denoted when the nose-point of the mouse was within the 2 cm^2^ border around each object to determine how much time was spent interacting with the objects. A discrimination index was calculated (Novel-Familiar)/Familiar) to determine if the mice spent more time with their familiar vs. their novel object. Mice were singly housed and acclimated to the testing area with a white noise machine for 1 hour prior to testing for both the open field and Novel Object tests.

### Tissue Collection

After euthanasia, the heart, liver, spleen, soleus, gastrocnemius, and white adipose tissue were collected, and their wet weights were recorded. The uterus of the female mice was also collected and weighed. In addition, the carotid artery, cerebral cortex, cerebral arteries (combination of middle cerebral artery, anterior cerebral arteries, and basilar arteries), and hippocampus were excised and flash frozen in liquid nitrogen for preservation. The right hemisphere of the brain was preserved in 4% paraformaldehyde for two days post-dissection and stored in 1x PBS solution and paraffinized until cryosectioning.

### Vascular Reactivity

Following euthanasia of the mouse, the brain was retrieved via decapitation, and the posterior cerebral arteries (PCAs) were dissected out of the cortex. PCAs were cannulated onto glass micropipette tips in a myograph chamber (Danish Myo Technology, Hinnerup, Denmark) filled with 145 mM NaCl, 4.7 mM KCl, 2 mM CaCl_2_, 1.17mMMgSO_4_, 1.2mM NaH2PO_4_, 5.0mM glucose,2.0mM pyruvate, 0.02mM EDTA, 3.0 mM MOPS buffer, 10 g/L BSA, 7.4 pH at 37C. The PCA was pressurized to 50mmHg over 20 minutes to mimic the internal pressure of the artery in vivo. Endothelial function was measured as a percent maximal or minimal luminal diameter following increasing concentrations of acetylcholine (ACh: 1×10^−9^ to 1×10^−4^ M). Endothelium-independent dilation was measured using sodium nitroprusside (SNP: 1×10^−10^ to 1×10^−4^ M). Before dose responses with ACh and SNP, the artery was pre-constricted using phenylephrine (1-6µM) to achieve >20% of the original luminal diameter. The PCA was incubated with endothelial nitric oxide inhibitor, N omega-nirto-L-arginine methyl ester (L-NAME, 30 minutes, 0.1mM). ACh doses were repeated after L-NAME incubation to quantify the NO-dependent dilation.

### Ex Vivo Pulse Pressure Application

PCA endothelial-dependent dilation in response to acute pulse pressure was assessed by *ex vivo* pulsation exposure. A separate PCA was cannulated into the pressure myograph chamber (Product #112PP, Danish Myo Technology, Hinnerup, Denmark). Pulsation at 400s-1 to mimicked a “low” pulse pressure (75mmHg to 50mmHg) and then a “high” pulse pressure (87.5 mmHg to 37.5 mmHg) for 30 minutes. The response to ACh was assessed at static pressure and again after the exposure to low and high pulsatile pressure. Since the exact PP in the PCA is unknown, the pressures were estimated based on the carotid artery PP in a mouse model of higher large artery stiffness^39^.

### Passive Stiffness, Diameter, Wall Thickness, and Wall: Lumen Ratio

Following endothelial-dependent dilation dose responses, the PCA and carotid arteries were placed in a calcium-free physiological salt solution for one hour. Then, the PCA experienced consecutive increases in pressure. To obtain β stiffness, stress was initially calculated as σ = PD/2WT, where P is the pressure in dynes/cm2, D is the lumen diameter, and WT is the wall thickness. Strain was further calculated as = (*D*-*D_i_*)/*D_i_*, where D*_i_* is the initial starting diameter. Data for each artery were fit to the curve using σ = σ_i_e^βε^ and β is the slope of the tangential elastic modulus versus stress. A higher β-stiffness indicates a stiffer artery ^40–42^. The Wall: lumen ratio was calculated as WT/*D* at 50 mmHg where WT is the wall thickness and *D* is the lumen diameter.

### Artery Histology

The middle cerebral artery (MCA) and a section of the carotid artery proximal to the carotid bifurcation was excised, dyed in Fast Green FCF, and flash frozen in optimal cutting temperature (OCT) compound for cyrosectioning at 8 um sections (Leica, Wetzlar, Germany) and adhered to a charged microscope slide. The MCA and carotid arteries were blocked with 20% goat serum for 15 min in a humidifier chamber, followed by two-hour incubation with anti-collagen-I or anti-collagen-III antibodies (ab270993 1:250, ab7778 1:250). The slides were further washed with PBS and incubated with an anti-rabbit secondary antibody (Fisher A-11070 1:1000). MCA and carotid elastin images were captured by the autofluorescence in the FIT-C channel. Slides were mounted with Prolong Gold with DAPI (Cat. No. P36935, Life Technologies). Three artery sections were prepared and stained per mouse/antibody/artery, plus one artery section with no primary antibody. The slides were imaged with a GE DeltaVision Ultra High Resolution Widefield Microscope with deconvolution. Elastin, collagen I, collagen III, and α-smooth muscle actin content were analyzed by the mean gray intensity using FIJI by ImageJ software. Values were averaged for the three artery sections from each mouse/antibody/artery, and the non-primary value was subtracted to obtain the final data values. The values were normalized to the male *E3xhAβPP* group.

### Immunohistochemistry

Paraffinized brains were cut at 5 µm thickness and incubated with primary antibodies for ionized calcium-binding adapter molecule 1 (Iba1) (ProteinTech, Rosemont, IL, USA, 10904-1-AP, 1:5000) or glial fibrillary acidic protein (GFAP) (ProteinTech, 16852-1-AP, 1:500), followed by development using Vector ELITE ABC kits (Vector Laboratories, Newark, CA, USA, PK-6102). Slices were stained with 3,3′-diaminobenzidine for visualization ^43^. Regions of interest (hippocampus, entorhinal cortex and thalamus) were determined using the Allen Brain Atlas and images were collected at a consistent area size (600 × 450 µm) using a Zeiss Axio Imager AZ10 (Zeiss Microscopy, Oberkochen, Germany) and analyzed using FIJI by ImageJ software (NIH, Bethesda, MD, USA) by percentage positive area. No positive staining for Aβ (4G8) was observed in any sample, and therefore, we were unable to quantify this marker.

### Gene Expression

mRNA from samples of the cortex and cerebral arteries (combined middle cerebral artery, anterior cerebral arteries, and basilar arteries) were quantified for expression of superoxide dismutase *(Sod)-1, Sod-2, Sod-3*, nicotinamide adenine dinucleotide phosphate oxidase2 (*Nox2*), and interleukin-1 β (*Il1b*) through qPCR. In samples of carotid arteries, mRNA gene expression was quantified for platelet-derived growth factor receptor alpha (*Pdgfr*α) and transforming growth factor β (*Tgfβ*). RNA from the cortex, cerebral arteries, and carotid artery were obtained using out laboratory standard protocol^44^. Values were normalized to the male *E3xhAβPP* group. The primers used are organized in **Supplemental Table 1**.

### Protein Expression

The hippocampus was dissected out of the cortex and flash-frozen with liquid nitrogen in an Eppendorf tube. Samples were homogenized in protein-extracting buffer and protease inhibitor cocktail, then directly sonicated twice at 50% power for 60-s intervals. Hippocampal lysates were centrifuged at 3,000 rpm for 10 minutes at 4°C, and the supernatant was removed. Lysate protein concentration was assessed using a commercially available kit (Pierce BCA protein assay kit; Thermo Fisher, Waltham, MA). Bio-Rad stain-free TGX gels were used for protein separation (4-20% gradient). The gels were activated after electrophoresis to ensure protein separation before being transferred onto a nitrocellulose membrane. Post transfer, the membranes were blocked with 5% bovine serum albumin (BSA) in TBS-T for 30 minutes, then incubated in primary antibodies overnight in PBS containing 3% BSA and 0.05% sodium azide. The primary antibody concentration for APOE was 1:1,000. Following incubation, membranes were washed and incubated in anti-rabbit horseradish peroxidase (HRP)-conjugated secondary antibody for 45 minutes. Following washes with TBS-triton, membranes were rinsed with the sensitivity enhancing Bio-Rad reagent, Clarity Western ECL Substrate, and imaged with a ChemiDoc Imaging System (ChemiDoc, Bio-Rad, Hercules, CA). Bio-Rad Image Lab software was used to determine protein band density, and relative protein expression was calculated as protein density normalized to total protein density. Values were normalized to the male *E3xhAβPP* group. See **Supplemental Table 2** for antibody list.

### Statistical Analysis

Statistical analyses were completed using SPSS 26.0 software, and GraphPad Prism 10.0. All data were examined for normal distribution according to Shapiro-Wilks tests and outliers with a z score of 2 or above were removed from analyses. Non-normally distributed data were transformed to achieve normality. A two-way ANOVA between groups was used for all outcomes except vascular reactivity data which was analyzed using a repeated measures ANOVA. Significance was set at p<0.05 and values were represented as mean ± SEM. Outliers were identified as those with *z*-score>2 and were removed from the dataset.

## Results

### Animal Characteristics

There was a main effect of both sex and genotype for body mass, with *E4/hAβPP* and male mice having a higher mass (**Supplemental Table 2**). *E3/hAβPP* males had significantly higher liver, percent liver mass normalized to body mass, and gastrocnemius mass than the *E3/hAβPP* females. *E4/hAβPP* males had significantly higher percent liver mass normalized to body mass than the *E4/hAβPP* females. *E3/hAβPP* females had significantly higher percent soleus normalized to body mass, uterus mass, and percent uterus normalized to body mass than *E4/hAβPP* females (**Supplemental Table 2**).

### Cognitive testing

Nest-building was used to assess instinctive behavior. There was a main effect of genotype, with *E4/hAβPP* mice having lower scores than *E3/hAβPP* mice **(Fig 1A)**. There were no differences in sex or genotype for the time spent on the accelerating rotarod **(Fig 1B)**. Similarly, there were no differences in time spent in the center of the open field test, indicating that the mice did not exhibit differences in anxious behaviors **(Fig 1C)**. However, there was an interaction of genotype and sex for the velocity and distance moved during the open field test, with the female *E4/hAβPP* mice being highest **(Fig 1D-E)**, suggesting that *E4/hAβPP* mice exhibit hyperactivity-like behaviors. The Novel Object Recognition was used to assess short-term and spatial memory. There was a main effect of genotype, with the *E4/hAβPP* mice spending less time with their novel object than the *E3/hAβPP* **(Fig 1F)**, suggesting impaired memory in the *E4/hAβPP* mice. Overall, the cognitive testing results indicate that the *E4/hAβPP* genotype has a negative impact on instinctual behavior and spatial memory, compared with the *E3/hAβPP* genotype.

**Figure 1.**
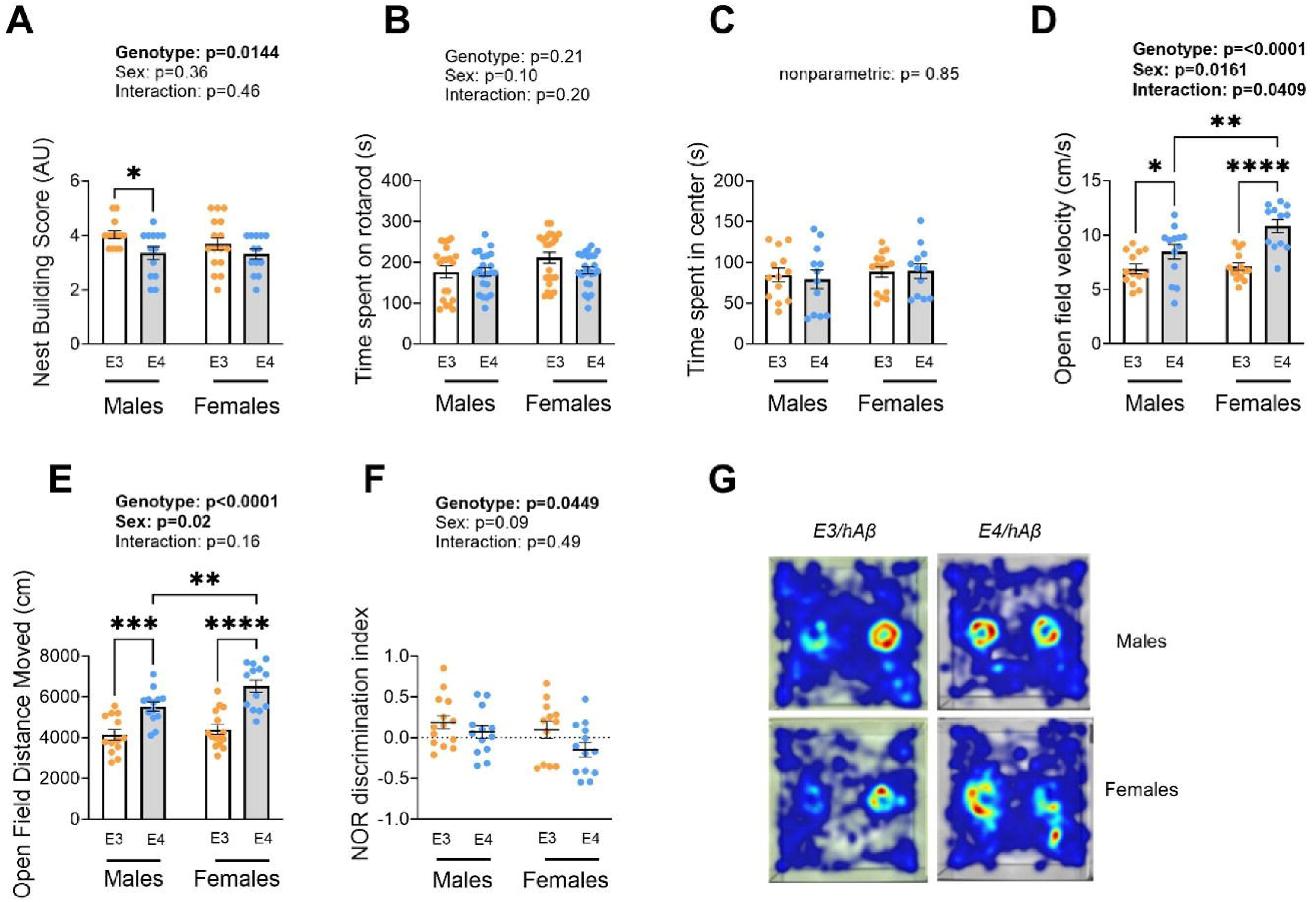
Instinctual behavior and memory are impaired in the *E4*x*hAβPP* mice. In male and female *E3*x*hAβPP* and *E4*x*hAβPP* mice, nest building test scores (*A)*, time spent on the accelerating rotarod (*B*), time spent in the center of the open field (*C*), velocity (cm/s) during the open field (*D*), distance (cm) moved during the open field test, and the discrimination index of time spent with the novel object (*F*). Representative heat maps of group averages for the novel object recognition test (*G*). A two-way ANOVA with Tukey’s multiple comparisons was used except for the open field time spent in the center where non-parametric t-tests were conducted between groups due to lack of normality. Data are mean ± SEM.

### Neuroinflammation and Neuropathology

Activated microglia and astrocytes were visualized by IHC to assess neuroinflammation. No detectable Aβ plaques were observed in either genotype (data not shown). There was a main effect of genotype, with the *E4/hAβPP* mice having more coverage of activated microglia (IBA1) in the hippocampus compared to the *E3/hAβPP* genotype, and with male *E4/hAβPP* having the highest coverage of IBA1 **(Fig 2A)**. There was a main effect of sex, with males having higher IBA1 coverage in the thalamus, but no effect of genotype **(Fig 2B)**. There were no differences observed between sexes or genotypes for the coverage of IBA1 in the entorhinal cortex **(Fig 2C)**. We assessed the same three brain regions, hippocampus, thalamus, and entorhinal cortex, for activated astrocytes (GFAP). There were no sex or genotypic differences observed in GFAP coverage across any of the three brain regions **(Fig 2E-G)**. Together, our results indicate that the *E4/hAβPP* genotype and male sex contribute to the accumulation of more activated microglia than activated astrocytes in the brain.

**Figure 2.**
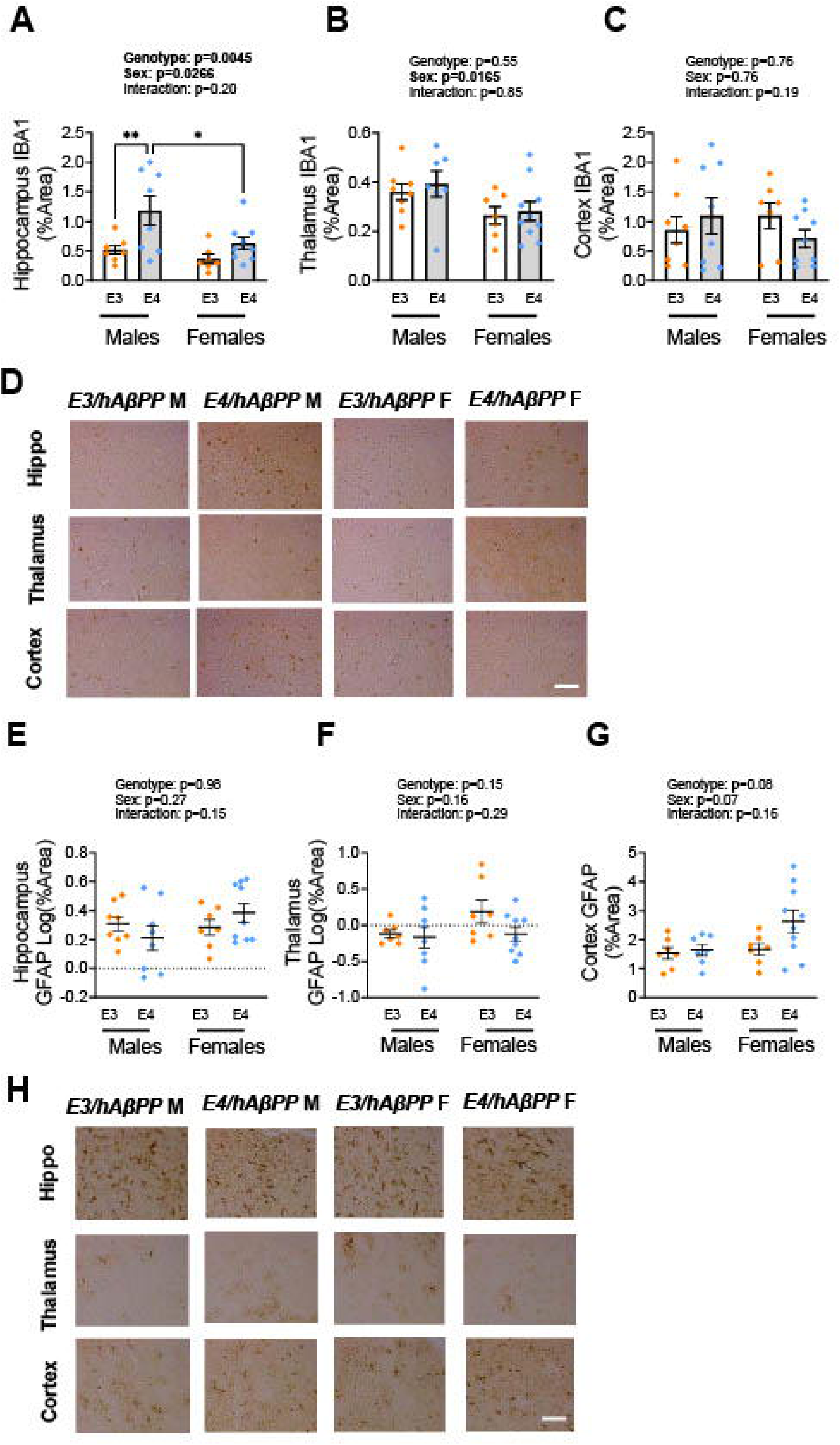
Male sex and *E4*x*hAβPP* genotype are associated with a higher activated microglia content. Paraffin-embedded slices from male and female *E4*x*hAβPP* and *E3*x*hAβPP* mice underwent IHC for activated microglia (IBA1) in the hippocampus (*A*), thalamus (*B*), and entorhinal cortex (*C*). Representative images of IBA1(*D*). The same groups were also probed for activated astrocytes (GFAP) in the hippocampus (E), thalamus (*F*), and entorhinal cortex (*G*). Representative images of GFAP (*H*). Scale bar: 200 µm. *n* = 6-9/group. *P < 0.05; **P <0.01. A two-way ANOVA with Tukey’s multiple comparisons was used. GFAP hippocampus and cortex data were Log transformed to achieve normality. Data are mean ± SEM.

### Large artery stiffness

Stiffer arteries have a reduced ability to dampen pulse pressure, thereby allowing higher pulse pressure to reach the smaller vessels in the brain, leading to dysfunction^45–47^. Therefore, we assessed the structure and stiffness of large elastic arteries. When we assessed the structure of the carotid arteries, we found the *E4/hAβPP* genotype carotids had a greater wall: lumen ratio than the *E3/hAβPP* genotype carotids (**Fig 3A**). However, this genotype effect was not statistically significant for carotid wall thickness alone (**Fig 3B**). As a result of higher collagen content and elastin fragmentation, the large arteries become stiffer. Thus, we assessed these parameters with immunofluorescence in the carotid arteries and stiffness in the aorta and carotid arteries. We found no genotype differences in elastin or collagen III content, but there was an effect of sex on collagen I content, with males having higher collagen I content than females (**Fig 3 C-E**). As a result, we find that *E4/hAβPP* mice have thicker carotid artery walls, but this thickness is not explained by differences in the extracellular matrix components we measured. We found no effect of genotype on aortic stiffness, measured by the gold standard method of pulse wave velocity^48^, but there was an effect of sex with *E3/hAβPP* males having higher aortic stiffness than *E3/hAβPP* females (**Fig 3F**). For the assessment of passive stiffness, the carotids were incubated in a Ca2+-free solution to eliminate the effect of smooth muscle tone. After incubation, the carotids underwent increasing luminal pressures to calculate the stress-strain curves (**Fig 3H**). From the stress-strain curves, we calculated the β stiffness and found the *E4/hAβPP* mice to have stiffer carotid arteries than the *E3/hAβPP* mice (**Fig 3I**). Overall, while the *E4/hAβPP* mice exhibit greater carotid stiffness and a higher wall: lumen ratio, the differences observed do not seem to be a result of extracellular matrix remodeling.

**Figure 3.**
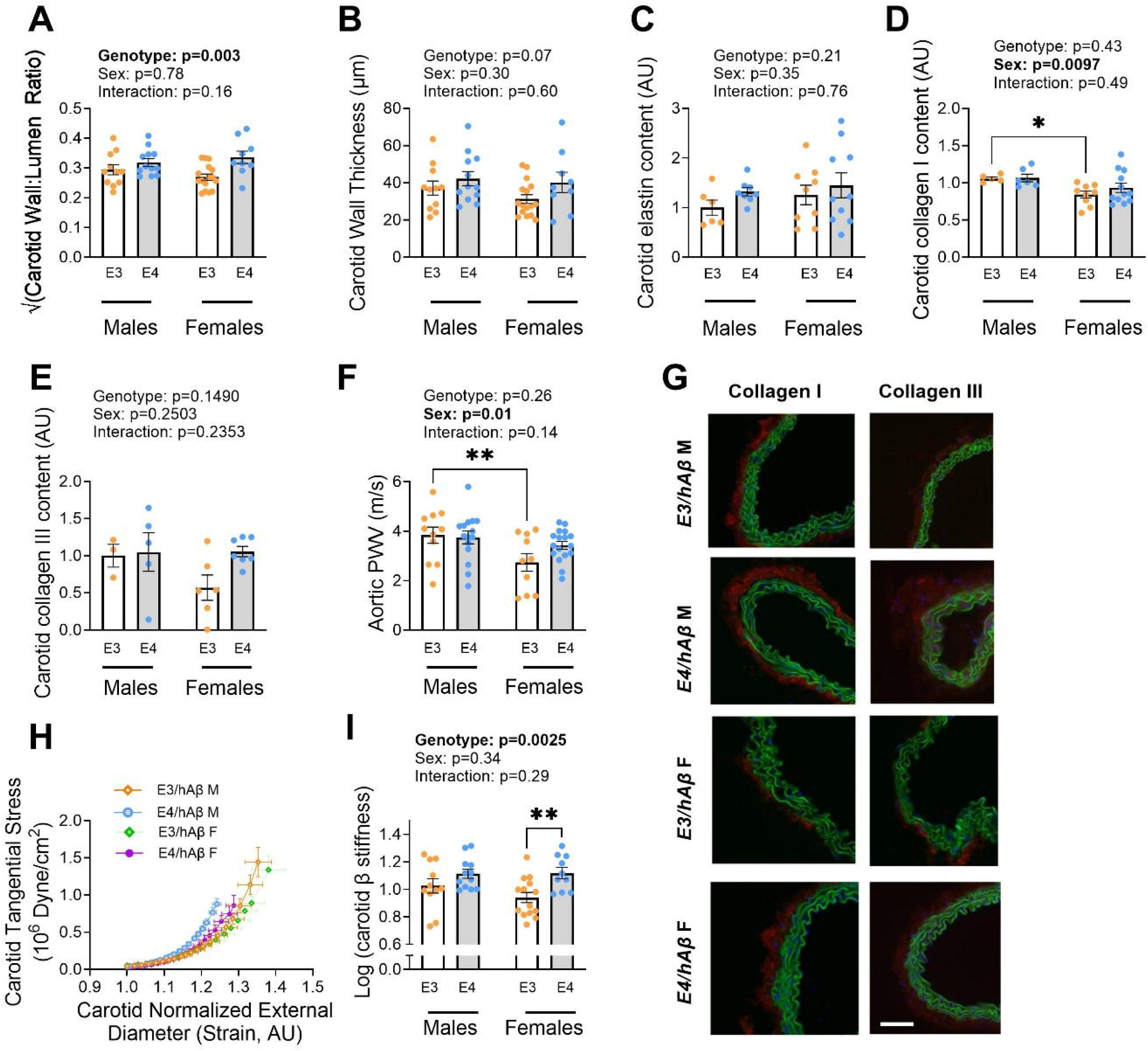
*E4*x*hAβPP* genotype is associated with higher carotid wall: lumen ratio and β stiffness. Carotid wall: lumen ratio (*A*) and wall thickness (*B*) measured by pressure myography in male and female *E4*x*hAβPP* and *E3*x*hAβPP* mice. Carotid elastin content (*C*), collagen I content (*D*), and collagen III content (*E*) measured by immunofluorescence in the male and female *E4*x*hAβPP* and *E3*x*hAβPP* mice. Aortic stiffness measured by PWV (*F*). Carotid immunofluorescence representative images: blue, DAPI; green, elastin; red, protein of interest (*G*). After incubation in a calcium-free solution, stress-strain curves were graphed for the passive stiffness (*H*) and the carotid β stiffness was measured in male and female *E4*x*hAβPP* and *E3*x*hAβPP* mice (*I*). Scale bar: 100 µm. *n* = 3-14/group. *P < 0.05; **P <0.01. A two-way ANOVA with Tukey’s multiple comparisons was used. The wall: lumen ratio was sqrt transformed and the β stiffness data was Log transformed to achieve normality. Data are mean ± SEM.

### Cerebral artery stiffness

As a result of large artery stiffness and less damping of high pulse pressures in the brain, the cerebral arteries will remodel and stiffen^24,49,50^. Thus, we also assessed the structure and stiffness of the PCA. Although there was a trend for the effect of sex and genotype on the PCA wall: lumen ratio and wall thickness, there were no statistically significant differences (**Fig 4A-B**). Given that stiffening of the cerebral arteries would impact the ability to dampen higher pulsatile pressures, we also assessed elastin and collagen content of the middle cerebral arteries. Similarly, to the carotid artery, we found no genotype differences in elastin or collagen I content; however, the *E4/hAβPP* mice had higher collagen I content than the *E3/hAβPP* mice (**Fig 4C-E**). We also assessed the passive stiffness of the PCA by generating stress-strain curves (Fig. **4G**). From our stress-strain curves, we found the *E4/hAβPP* mice had higher PCA β stiffness than the *E3/hAβPP* mice (**Fig 4H**). Overall, our findings suggest the *E4/hAβPP* genotype impacts cerebral artery structure, resulting in higher cerebral artery stiffness.

**Figure 4.**
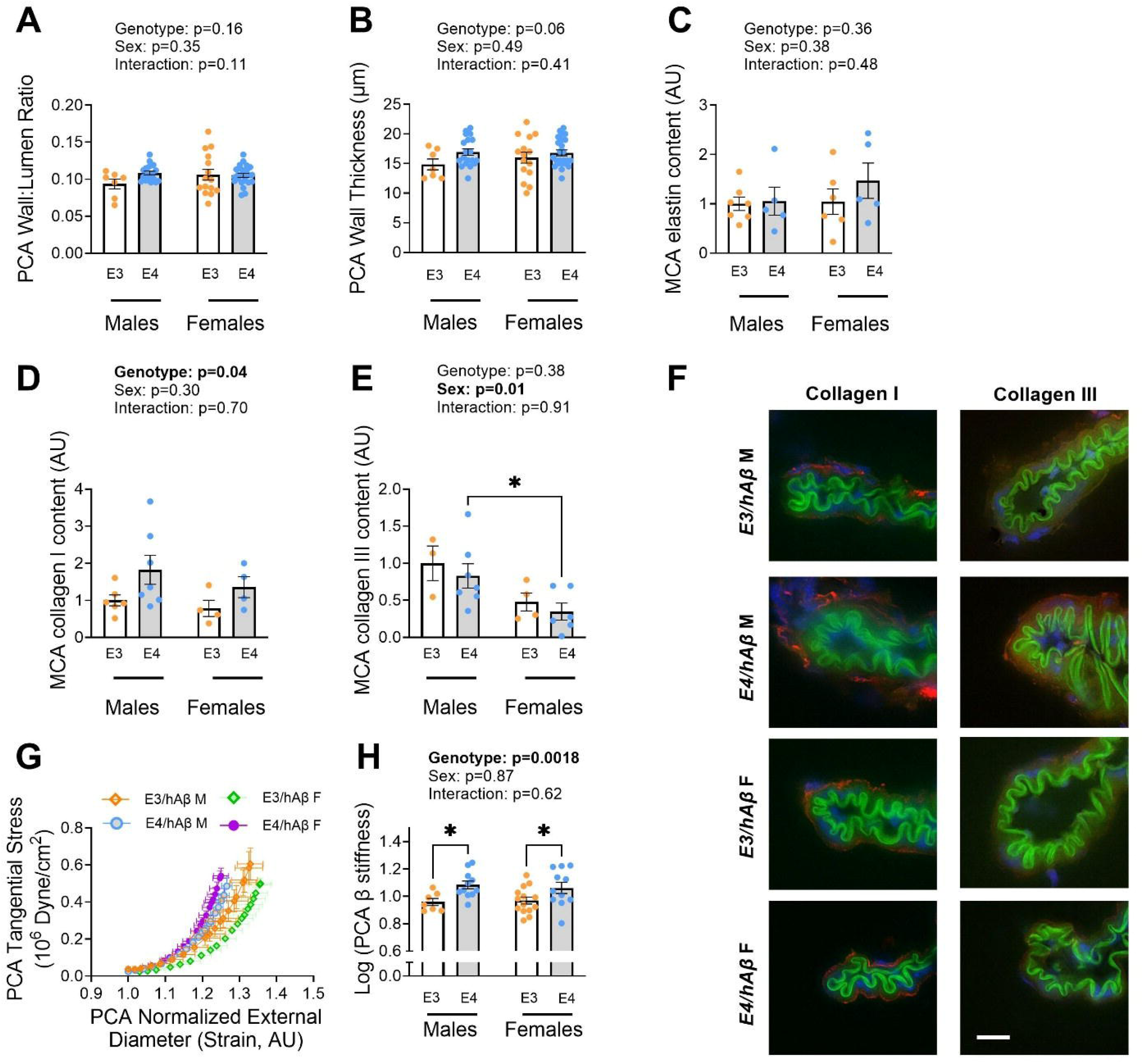
*E4*x*hAβPP* have greater cerebral artery stiffness. Posterior cerebral artery (PCA) wall: lumen ratio (*A*) and wall thickness (*B*) measured by pressure myography in male and female *E4*x*hAβPP* and *E3*x*hAβPP* mice. Middle cerebral artery (MCA) elastin content (*C*), collagen I content (*D*), and collagen III content (*E*), measured by immunofluorescence in the male and female *E4*x*hAβPP* and *E3*x*hAβPP* mice. Representative images: blue, DAPI; green, elastin; red, protein of interest (*F*). After PCA incubation in a calcium-free solution, the stress strain curves (*G*), and β stiffness (*H*) were measured by pressure myography in the male and female *E3*x*hAβPP* and *E4*x*hAβPP* mice. The PCA β stiffness data was Log transformed to achieve normality. A two-way ANOVA with Tukey’s multiple comparisons was used. *P < 0.05 Data are mean ± SEM.

### Vascular function

Previous studies have yet to explore the *E4/hAβPP* genotype effects on cerebrovascular function. We found no genotype or sex differences in the PCA vasodilation in response to ACh, which was confirmed to be nitric oxide synthase mediated because of the little to no vasodilation after the LNAME incubation (**Fig 5A**). Similarly, we found no group differences in endothelium-independent dilation in response to SNP (**Fig 5B**). Thus, we saw no impact of *APOE* genotype on cerebral artery function. However, arteries are not exposed to static pressure *in vivo*; rather, they experience pulsatile pressure. Elevated pulsatile pressure is known to induce endothelial cell dysfunction^51,52^. Thus, we assessed the interaction effect of the *APOE* genotype and pulse pressure on cerebral artery endothelium-dependent vasodilation by exposing the PCA *ex vivo* to static, low pulsatile, and high pulsatile pressures. We found no impairments in PCA vasodilation after low pulse pressure, but high pulse pressure significantly impaired PCA vasodilation in response to ACh in the *E4/hAβPP* mice but not in the *E3/hAβPP* mice (**Fig 6A-D**). Furthermore, for the high pulse pressure condition, maximal vasodilation to ACh was lower in the *E4/hAβPP* female mice compared with the *E3/hAβPP* female mice (**Fig 6E**). There was no effect of sex or genotype on spontaneous tone of the PCA during pulsing conditions (**Supplemental Table 3**). Together, our vascular function results suggest that high pulse pressure is more detrimental to the *E4/hAβPP* genotype than to the *E3/hAβPP* genotype, with the greatest effect in *E4/hAβPP* females.

**Figure 5.**
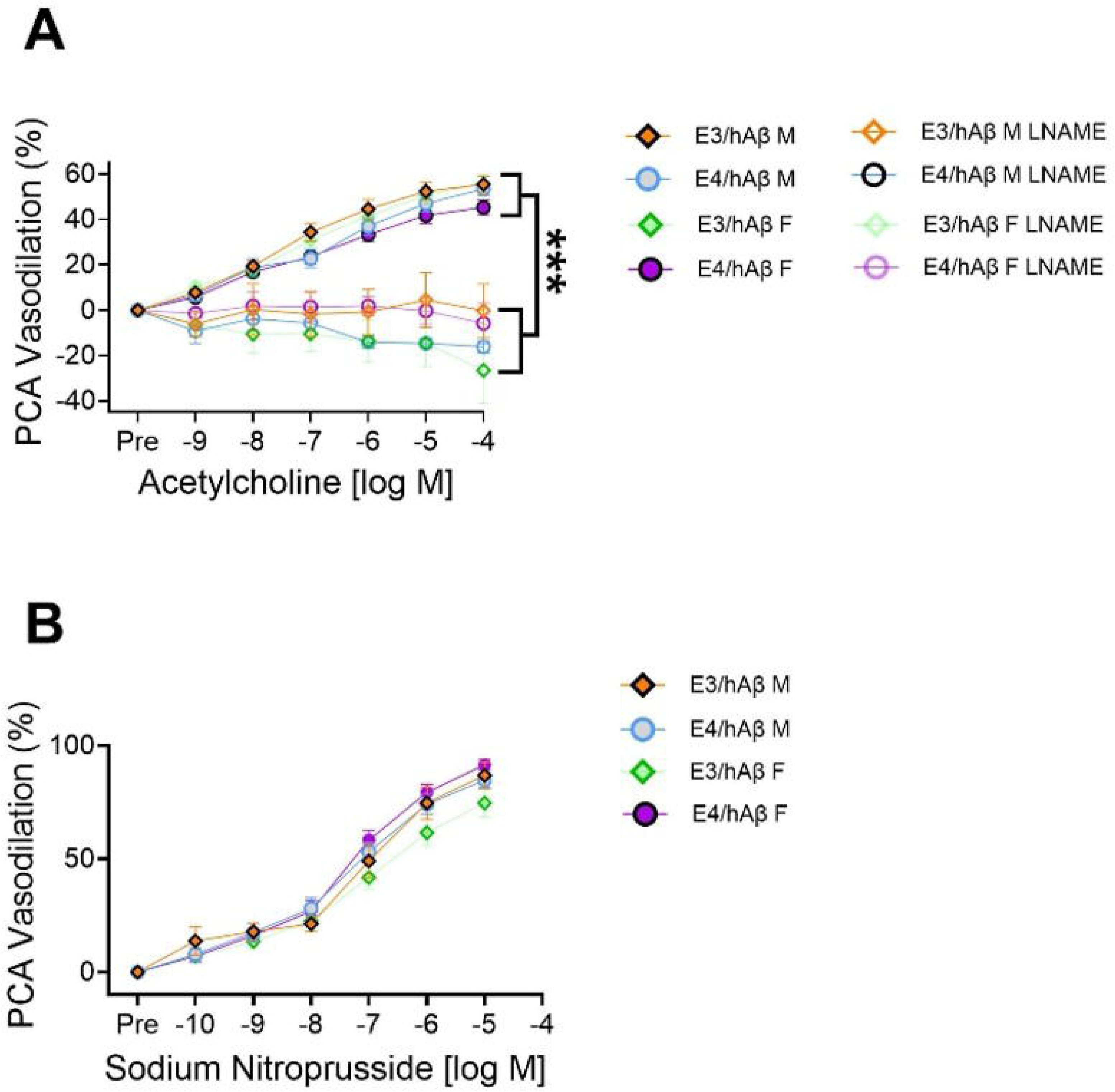
Sex and genotype do not affect cerebral artery endothelium-dependent or endothelium-independent vasodilation during static pressure. Posterior cerebral artery (PCA) endothelium-dependent dilation to acetylcholine with and without the presence of nitric oxide synthase inhibitor, LNAME, (*A*) and endothelium-independent dilation to sodium nitroprusside (B) in *E3*x*hAβPP* and *E4*x*hAβPP* males and females. *n*=4-15/group. ***P < 0.001. A repeated measures ANOVA with Tukey’s multiple comparisons was used. Data are mean ± SEM.

**Figure 6.**
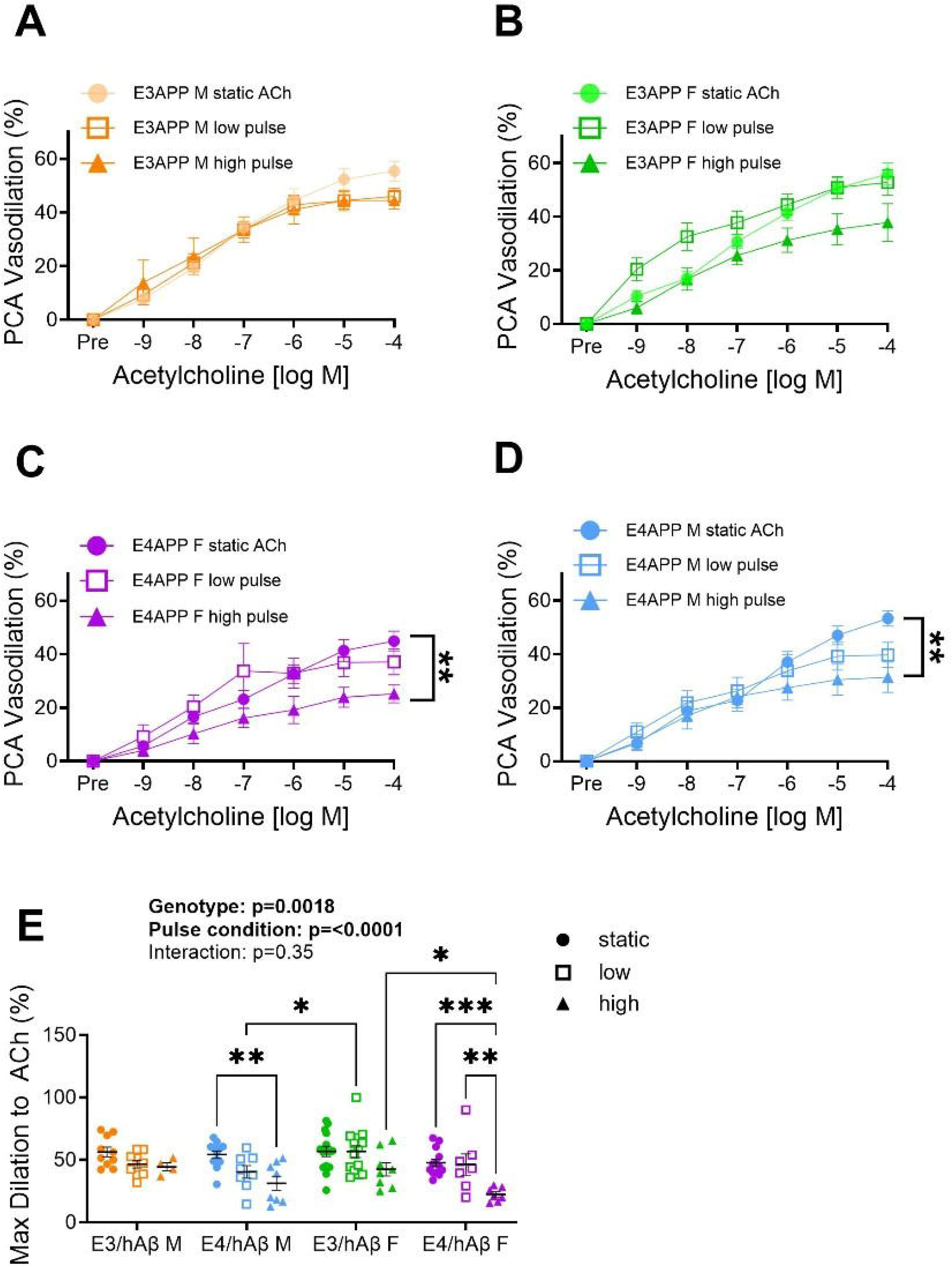
*E4*x*hAβPP* genotype was most susceptible to the negative impact of high pulse pressure on cerebral artery vasodilation. Posterior cerebral artery (PCA) endothelium-dependent dilation in *E3xhAβPP* males (*A*), *E3xhAβPP* females (*B*), *E4*xhAβPP females (*C*), and *E4*xhAβPP males (*D*) in response to acetylcholine after exposure to static pressure, low pulse pressure, and high pulse pressures. Maximal PCA vasodilation to acetylcholine in both male and female *E3*xhAβPP and *E4*xhAβPP mice following static, low, and high pulse pressures (E). *n* = 4-15/group. *P < 0.05; **P <0.01. A repeated measures and two-way ANOVA with Tukey’s multiple comparisons was used. Data are mean ± SEM.

### Markers of oxidative stress, inflammation, and vascular remodeling

To determine potential mechanisms for the effects of the *E4/hAβPP* genotype on vascular structure and function and neuroinflammation, we assessed oxidative stress and pro-inflammatory markers in the cortex, cerebral arteries, and carotid arteries. We observed that the *E4/hAβPP* genotype had higher *Nox2* and *Sod1* expression in the cortex compared to the *E3/hAβPP* genotype (**Fig 7A-B**). Alternatively, we found no significant effects of sex or genotype on *Sod2, Sod3, or Il1b* in the cortex (**Fig 7C-E**). Our cortex gene expression results suggest heightened pro-oxidant activity in the *E4/hAβPP* genotype, which may cause a compensatory increase in the antioxidant *Sod1*. In the cerebral arteries, while there was no effect of genotype on the expression of *Nox2*, there was a main effect of sex, with females having higher *Nox2* expression compared to the males (**Fig 7F**). Our results indicated no group differences in cerebral artery *Sods 1, 2,* or *3* expression (**Fig 7G-I**). Interestingly, we observed that the *E4/hAβPP* genotype had higher *Il1b* expression in the cerebral arteries than *E3/hAβPP* (**Fig 7J**). Higher expression of *Il1b* in the cerebral arteries, but not the cortex, may be due to a greater effect of the *E4/hAβPP* genotype on cerebral artery inflammation than on cortical inflammation. Lastly, we assessed *Pdgfr*α and *Tgfβ*, which are involved in arterial remodeling signaling^53^. We observed no genotype or sex differences in *Pdgfr*α nor *Tgfβ* in the carotid artery (**Fig 7K-L**). To determine if the *E4/hAβPP* genotype influences APOE expression, we measured APOE protein expression in the hippocampus. We observed no genotypical or sex differences in hippocampal APOE expression **(Fig 7M)**. Thus, the overall expression of the APOE protein may not play a critical role in modulating the effects of the *E4/hAβPP* genotype. Our gene expression and protein results suggest that *E4/hAβPP* mice have a pro-oxidative environment in the cortex and a pro-inflammatory environment in the cerebral arteries.

**Figure 7.**
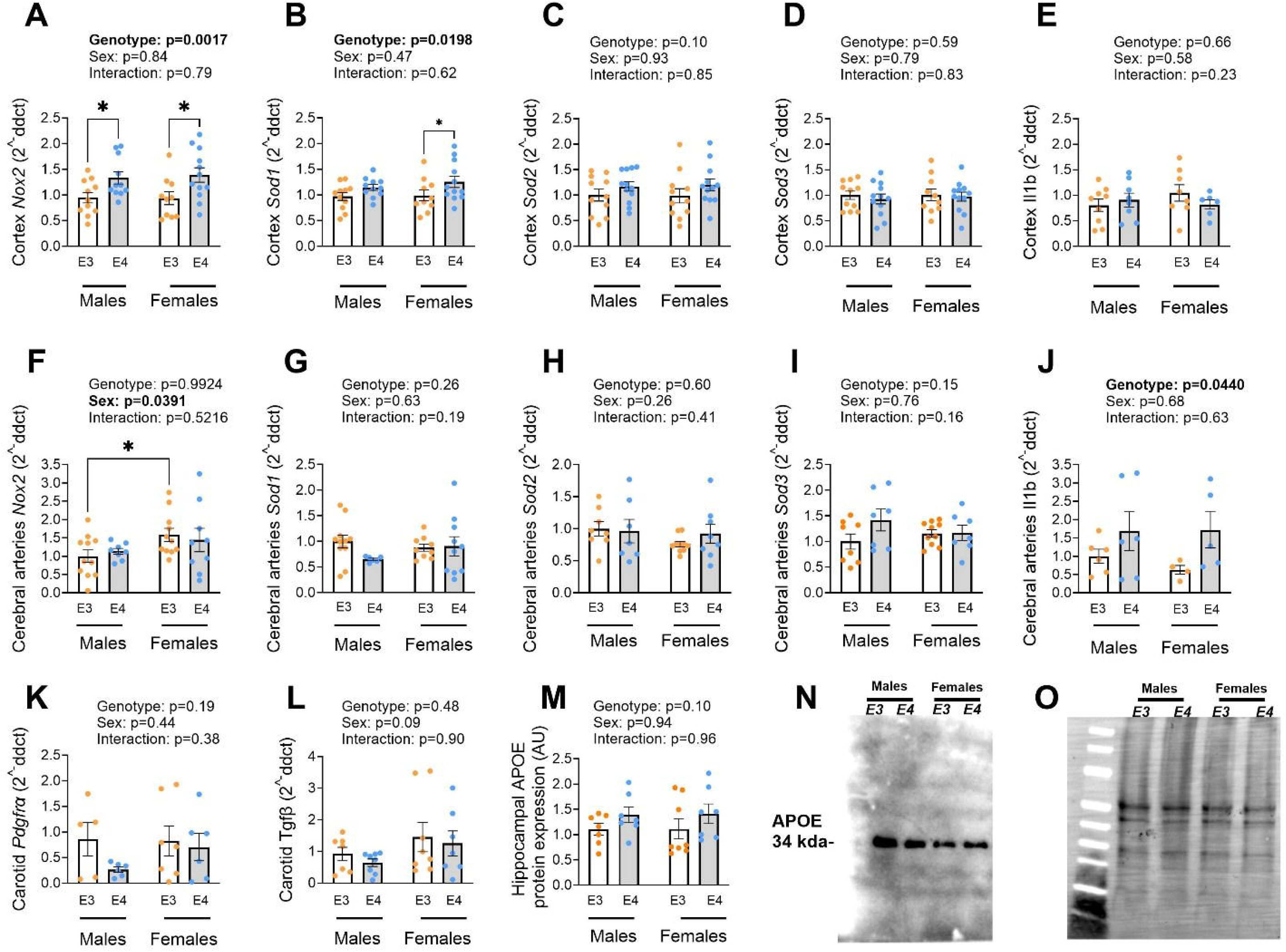
The *E4*x*hAβPP* genotype exhibited higher markers of oxidative stress in the cortex and inflammation in the cerebral arteries. Cortex and cerebral artery gene expression of *Nox2* (nicotinamide adenine dinucleotide phosphate oxidase 2) (*A,F*), *Sod1* (superoxide dismutase 1) (*B,G*), *Sod2* (*C,H*), *Sod3* (*D,I*), and *Il1b* (interleukin-1β) (*E,J*) in *E4*x*hAβPP* and *E3*x*hAβPP* male and females. Carotid artery gene expression of *Pdgfr*α (platelet-derived growth factor receptor α) (*K*) and *Tgfβ* (transforming growth factor β) (*L*) in *E4*x*hAβPP* and *E3*x*hAβPP* male and female mice. Hippocampal APOE protein expression (*M*), representative blot image of protein of interest, APOE (*N*) and total protein (*O*). *n* = 5-10/group. *P < 0.05. *Nox2* and *Sod1* gene expression data were transformed by sqrt to achieve normality. A two-way ANOVA with Tukey’s multiple comparisons was used. Data are mean ± SEM.

## Discussion

In this study, we demonstrate the utility of the *E4/hAβPP* mouse model as a more translational approach to understanding how the *E4* genotype increases AD risk. The *E4/hAβPP* mouse has a multifaceted phenotype characterized by cognitive impairment, neuroinflammation, vascular remodeling, and heightened susceptibility to cerebrovascular stress. Collectively, our findings support the concept that the *E4* genotype has influences beyond amyloid-related pathology by contributing to the vascular and inflammatory processes increasingly recognized as critical drivers of AD progression.

### APOE4 promotes cognitive impairment and region-specific neuroinflammation

The *E4/hAβPP* mice exhibited deficits in both nest-building and novel object recognition, indicating impairments in instinctual behavior and hippocampal-dependent memory. Our findings are consistent with a growing body of literature demonstrating that *E4* impairs cognition in mouse models^54^. Human *E4* targeted-replacement mice exhibit deficits in learning and memory tasks, including impaired spatial learning and memory in the Barnes Maze and novel object recognition task compared with *E3* mice, even in the absence of extensive AD pathology^17,55,56^. These findings suggest that *E4* may confer susceptibility to cognitive dysfunction even before substantial amyloid pathology develops.

Neuroinflammation is increasingly recognized as a central contributor to AD pathogenesis, and the present findings suggest that the *E4/hAβPP* mice promote region-specific microglial activation. We observed increased hippocampal IBA1 coverage in *E4/hAβPP* mice, with *E4/hAβPP* males exhibiting the greatest degree of microglial activation. Our findings align with literature from both human and mouse brain tissue demonstrating that the *E4* genotype influences microglial function and promotes inflammatory signaling ^57–59^. In contrast, no genotype differences were observed in the entorhinal cortex, suggesting that inflammatory responses may develop selectively across brain regions. The absence of differences in the entorhinal cortex is somewhat unexpected given the early vulnerability of this region in human AD and may reflect the relatively young age of the mice or the early stage of pathology present in this model^60,61^.

In contrast to the observed higher microglial content in *E4/hAβPP* mice, astrocyte coverage did not differ across groups. This finding suggests that microglial activation may be a more sensitive indicator of early neuroinflammatory alterations in the *E4/hAβPP* model, although additional studies using complementary markers of astrocyte function are warranted.

### Artery stiffness may contribute to vascular vulnerability in E4/hAβPP mice

In addition to cognitive and neuroinflammatory differences, *E4/hAβPP* mice exhibited evidence of vascular stiffening characterized by higher carotid and PCA stiffness. Arterial stiffening is increasingly recognized as a contributor to AD risk because it enhances transmission of pulsatile stress to the cerebral circulation and promotes cerebrovascular dysfunction and impaired waste clearance mechanisms^45,46,62–64^. Consistent with established sex differences in vascular aging, males exhibited greater aortic PWV than females^65^. Although genotype differences were not observed for aortic PWV, this measure is influenced by several physiological factors *in vivo*, including blood pressure^66,67^ and vascular smooth muscle tone^68^. Notably, *E4*-associated effects emerged more prominently in the carotid and cerebral vasculature, where β-stiffness was significantly elevated compared with *E3/hAβPP* mice. These findings suggest that *E4* contributes to vascular remodeling in arterial beds that may be particularly important for maintaining brain health^49,69^.

The mechanisms contributing to the greater arterial stiffness observed in *E4/hAβPP* mice may differ between vascular beds. In the carotid arteries, greater stiffness occurred despite minimal alterations in extracellular matrix composition, suggesting that factors beyond the structural components measured in this study contribute to the observed mechanical outcomes. In contrast, cerebral arteries from *E4/hAβPP* mice exhibited both higher stiffness and greater collagen I content. Since collagen I is a major load-bearing component of the arterial wall and is associated with reduced vascular compliance^70^, greater collagen deposition may contribute directly to the cerebrovascular stiffness observed in the *E4/hAβPP* mice. Together, these findings suggest that *E4* genotype promotes structural remodeling of the cerebral vasculature that may heighten the transmission of high pulsatile energy into the brain and enhance vulnerability to cerebrovascular dysfunction.

### E4 is more susceptible to pulse pressure-induced cerebrovascular dysfunction

Under static pressure conditions, cerebral artery endothelial- and smooth muscle-dependent vasodilatory responses were similar across groups, suggesting that *E4*-associated vascular deficits are not readily detectable under basal conditions. Consistent with previous findings from our laboratory, these results indicate that cerebrovascular dysfunction may emerge only when the vasculature is exposed to physiological stressors^71,72^. This interpretation is supported by evidence that *E4* promotes cerebrovascular vulnerability before overt functional deficits become apparent^16^. Indeed, our findings suggest that *E4* in the presence of *hAβPP* reduces the ability of cerebral arteries to tolerate the hemodynamic stress associated with elevated pulsatility. This observation aligns with growing evidence that *E4* carriers exhibit heightened vulnerability to vascular risk factors, including hypertension, arterial stiffness, and cerebral hypoperfusion^73,74^. Recent human studies have demonstrated that the relationship between vascular burden and cognitive performance is stronger in *E4* carriers than in non-carriers^73^ and that elevated pulse pressure is associated with reduced CBF in AD-vulnerable brain regions, particularly among *E4* carriers^75^. Furthermore, *E4* has been linked to impaired neurovascular regulation^16^, reduced vascular integrity^19,76,77^, and altered cerebrovascular reactivity^17,78^, suggesting that *E4-*associated vascular dysfunction may remain latent under resting conditions and become apparent only when the cerebral vasculature is challenged.

### Oxidative stress and inflammation are elevated in E4/hAβPP mice

The *E4* genotype has previously been linked to increased oxidative stress, vascular dysfunction, and chronic neuroinflammatory signaling, all of which are thought to contribute to AD progression^55,59,79–81^. Our findings support heightened *E4*-dependent oxidative stress and inflammation as demonstrated by the higher expression of *Nox2* and *Sod1* in the cortex of *E4/hAβPP* mice. Since NADPH oxidase is a major source of reactive oxygen species in the brain and vasculature, higher *Nox2* expression may indicate elevated oxidative stress, whereas higher *Sod1* expression may represent a compensatory antioxidant response. Collectively, these findings support a genotype-dependent shift toward a more pro-oxidative and pro-inflammatory environment in *E4/hAβPP* mice compared to *E3/hAβPP* mice.

The most prominent inflammatory finding was the elevated cerebral artery *Il1b* expression in *E4/hAβPP* mice. Although the present study cannot establish a causal role for IL-1*β*, prior studies have demonstrated that perturbed IL-1β signaling leads to vascular remodeling and inflammatory vascular pathology, suggesting that elevated *Il1b* expression may contribute to the vascular phenotype observed in the *E4/hAβPP* mice^82^. Furthermore, studies utilizing humanized *E4* mouse models have demonstrated exaggerated inflammatory responses and dysfunctional immune signaling, supporting the biological plausibility of this interpretation^59,83^.

### Limitations

Several limitations should be considered when interpreting these findings. First, all data were collected in relatively young mice. Consequently, additional *E4*-associated pathological features that emerge with aging, including greater amyloid burden and more extensive neuroinflammation, may not yet be fully developed. Second, vascular function was assessed ex vivo and therefore does not fully recapitulate the complex hemodynamic environment present in vivo. While *E4/hAβPP* mice exhibited greater arterial stiffness and impaired vasodilatory responses following pulse pressure exposure, we did not measure cerebral blood flow and neurovascular coupling directly. Finally, the present study cannot establish causal relationships among oxidative stress, inflammation, vascular dysfunction, and cognitive impairment. Future studies incorporating longitudinal assessments and mechanistic interventions will be needed to determine how these processes interact during disease progression.

### Conclusions

Our findings are the first to show that *E4/hAβPP* mice exhibit convergent cognitive, inflammatory, and vascular abnormalities that are consistent with multiple features of AD. Most notably, high pulse pressure exposed a cerebrovascular vulnerability that was largely absent under baseline conditions in *E4/hAβPP mice*, suggesting that vascular stress may be a critical modifier of *E4*-associated risk. Together, our findings support a model in which *E4* amplifies susceptibility to vascular dysfunction, inflammation, and arterial stiffening, thereby enhancing vulnerability to cognitive decline. These results further establish the *APOE/hAβPP* model as a useful platform for investigating interactions among *E4* genotype, vascular dysfunction, and AD pathogenesis.

