## Supplemental data for "A humanized Aβ mouse model reveals *E4*-dependent cognitive impairments, microglial activation, and cerebrovascular dysfunction"

**Data Supplement**

Ashley E. Walker, PhD

Department of Human Physiology

University of Oregon

139 Straub Hall 1240 University of Oregon

Eugene, Oregon, 97403, USA

https://vascularlab.uoregon.edu/

**Supplemental Table 1. Gene Expression Primer Sequence**

| Primer | Forward | Reverse |
|---|---|---|
| SOD1 | AACCAGTTGTGTTGT CAG GAC | CCACCATGTTTCTTAGAGTGAGG |
| SOD2 | CAGACCTGCCTTACGACTATGG | CTCGGTGGCGTTGAGATTGTT |
| SOD3 | CCTTCTTGTTCTACGGCTTGC | TCGCCTATCTTCTCAACCAGG |
| PDGFRA | AGAGTTACACGTTTGAGCTGTC | GTCCCTCCACGGTACTCCT |
| MMP9 | GCAGAGGCATACTTGTACCG | TGATGTTATGATGGTCCCACTTG |
| NOX2 | TCCCAGAGAACACAGCATAAC | CTAGCCTGCTTATGGGATTCTT |
| IL1B | GCAACTGTTCCTGAACTCAACT | ATCTTTTGGGGTCCGTCAACT |
| TGF-B | CTCCCGTGGCTTCTAGTGC | GCCTTAGTTTGGACAGGATCTG |
| 18s | TAGAGGGACAAGTGGCGTTC | CGCTGAGCCAGTCAGTGT |

**Supplemental Table 2: Animal Characteristics**

| Variable | ***E3xhAβPP* Males** | ***E4xhAβPP* Males** | ***E3xhAβPP* Females** | ***E4xhβAPP* Females** |
|---|---|---|---|---|
| N | 11 | 9 | 13 | 14 |
| Age, months | 6±0.1 | 6±0.1 | 6±0.1 | 6±0.1 |
| Body mass (g)[a,b] | 29.39 ± 2.08 | 31.78 ± 0.84* | 21.76 ±1.23* | 23.76±2.69*†‡ |
| Heart mass (mg) | 0.15 ±0.02 | 0.15±0.02 | 0.13±0.09 | 0.14±0.08 |
| Percent heart:body mass | 0.50±0.07 | 0.48±0.05 | 0.60±0.36 | 0.60±0.41 |
| Liver mass (mg)[b] | 1.66 ±0.24 | 1.65 ±0.12 | 1.03 ±0.11* | 1.14±0.16† |
| Percent liver:body mass[b] | 5.65±0.65 | 5.18±0.35 | 4.75±0.58* | 4.82±0.60 |
| WAT mass (g) | 0.40±0.11 | 0.81±0.14 | 0.29±0.07 | 0.55±0.33 |
| Percent WAT:body mass | 1.34±0.34 | 2.53±0.40 | 1.35±0.33 | 2.21±1.12 |
| Spleen mass (mg) | 0.17±0.22 | 0.10±0.02 | 0.09±0.01 | 0.10±0.02 |
| Percent spleen:body mass | 0.60±0.85 | 0.32±0.49 | 0.41±0.05 | 0.42±0.08 |
| Gastroc mass (mg)[b] | 0.18±0.05 | 0.17±0.03 | 0.13±0.03* | 0.12±0.02 |
| Percent gastroc:body mass[a] | 0.62±0.16 | 0.52±0.11 | 0.59±0.14 | 0.53±0.1 |
| Soleus mass (mg) | 0.01±0.01 | 0.01±0.003 | 0.01±0.004 | 0.01±0.002 |
| Percent soleus:body mass[a] | 0.04±0.02 | 0.03±0.01 | 0.04±0.02 | 0.03±0.01‡ |
| Uterus mass (mg)[a] | | | 0.11±0.02 | 0.09±0.02 ‡ |
| Percent uterus:body mass[a] | | | 0.49±0.10 | 0.37±0.09 ‡ |

Data are mean ± SD. White adipose tissue (WAT), gastrocnemius (gastroc). [a] $p<0.05$ main effect of genotype, [b] $p<0.05$ main effect of sex, [c] $p<0.05$ interaction genotype x sex, *P<0.05 vs. *E3xhAβPP* males, †P<0.05 vs *E4xhAβPP* males, ‡P<0.05 vs *E3xhAβPP* females

**Supplemental Table 3: Posterior cerebral artery characteristics**

| Variable | *E3xhAβPP* Males | *E4xhAβPP* Males | *E3xhAβPP* Females | *E4xhβAPP* Females |
|---|---|---|---|---|
| **Spontaneous Tone (% maximal diameter)** | | | | |
| Static Pressure | 6.20±1.25 | 5.57±1.33 | 4.47±1.23 | 7.81±1.40 |
| Low PP | 6.39±2.08 | 5.85±2.65 | 5.90±1.19 | 13.75±2.99 |
| High PP | 5.22±2.81 | 3.14±1.30 | 4.43±1.63 | 9.07±2.55 |
| **Preconstriction Tone (% maximal diameter)** | | | | |
| Static Pressure | 30.57±1.33 | 35.97±2.23 | 34.31±1.98 | 36.87±1.93 |
| Low PP | 28.86±2.43 | 35.76±1.59 | 31.10±1.99 | 34.03±2.34 |
| High PP | 22.35±2.26 | 24.06±1.48 | 23.42±1.91 | 29.16±3.44 |

Data are mean ± SEM.
